# Force-dependent remodeling of cytoplasmic ZO-1 condensates contributes to robust cell-cell adhesion

**DOI:** 10.1101/2020.10.04.323436

**Authors:** Noriyuki Kinoshita, Takamasa S. Yamamoto, Naoko Yasue, Chiyo Takagi, Toshihiko Fujimori, Naoto Ueno

**Author notes:** corresponding authors (NU), (NK.).

## Abstract

Although the physiological importance of biomolecular condensates is widely recognized, how it is controlled in time and space during development is largely unknown. Here we show that a tight junction protein ZO-1 forms cytoplasmic condensates in the trophectoderm (TE) of the mouse embryo before E4.0. These disappear via dissolution, and ZO-1 accumulates at the cell junction as the blastocyst cavity grows and internal pressure on TE cells increases. In contrast, this dissolution was less evident in TE cells attached to the inner cell mass, as they receive weaker tensile forces. Furthermore, analyses using MDCK cells have demonstrated that the ZO-1 condensates are generated and maintained by liquid-liquid phase separation. Our study also highlights that the dynamics of these condensates depends on the physical environment via an interaction between ZO-1 and F-actin. We propose that the force-dependent regulation of ZO-1 condensation contributes to establishing robust cell-cell adhesion during early development.

## Introduction

Mechanical forces are generated at different times and in different tissues during the development of organisms due to dynamic movements and changes in the shape of cells and tissues (Bodor et al., 2020), growth of a cell mass by proliferation (Godard and Heisenberg, 2019), the removal of cells by apoptosis (Teng et al., 2017), luminal pressure (Chan et al., 2019), and shear stress of body fluids such as blood (Paolini and Abdelilah-Seyfried, 2018), etc. All of these forces can contribute to the morphogenesis of organs through mechanochemical feedback mechanisms (Hannezo and Heisenberg, 2019). Although the presence and physiological importance of such forces have long been implicated in the homeostasis as well as morphogenesis of living organisms (Hallou and Brunet, 2020; Thompson, 1917), how cells sense and respond to these forces is not fully understood at the molecular level. This study demonstrates that mechanical forces control the remodeling of ZO-1 through the regulation of liquid-liquid phase separation (LLPS), a physical process that facilitates the demixing of proteins through protein condensation.

In our previous study using *Xenopus laevis* embryos, we demonstrated that among a number of identified phosphoproteins, cell junction-as well as focal adhesion-related proteins are highly phosphorylated immediately after the application of mechanical forces by centrifugation or compression (Hashimoto et al., 2019). Intriguingly, a global analysis of the phosphoproteome suggested that mechanical stimuli induced the embryonic cells to adopt an epithelial state rather than a mesenchymal state, known as the mesenchymal-epithelial transition (MET), which is opposite to the epithelial-mesenchymal transition (EMT), a well-characterized phenomenon in some developmental contexts and in cancer pathogenesis in which cells compromise their cell-to-cell adhesiveness (Baum et al., 2008). In fact, we confirmed that the adherens and tight junction, revealed by the accumulation of C-cadherin (Choi et al., 1990; Ginsberg et al., 1991) and ZO-1 (Anderson et al., 1988), respectively, became enhanced after the application of force. We also found that F-actin, which is normally localized in the basal and lateral domain of early embryonic cells, accumulated at the apical domain, reinforcing the stiffness of the cell cortex (Hashimoto et al., 2019). These observations suggest that these junctions were remodeled following force stimuli and that the cells adopted more epithelialized states. This led us to propose that force-induced epithelialization is an important cellular response to physical forces to maintain tissue integrity, allowing it to acquire robustness against forces that might perturb normal morphogenesis.

More recently, we showed that FGFR1 is activated and that Erk2 is consequently phosphorylated and translocated into the nucleus in ectodermal cells (Kinoshita et al., 2020) to achieve MET during gastrulation. In that study, we repeatedly observed that ZO-1 condensates were found as cytoplasmic puncta in ectoderm cells prior to, but not after, gastrulation (Kinoshita et al., 2020). Based on previous reports that showed that ZO-1 could undergo LLPS in zebrafish embryos (Schwayer et al., 2019) and cultured cells (Beutel et al., 2019), we speculated that the ZO-1 LLPS is mechanically regulated, particularly by dynamic morphogenesis known as epiboly during gastrulation which imposes a tensile force on the ectoderm (Hernández-Vega et al., 2017).

To extend the above study that employed *Xenopus laevis* and to examine whether our working model could be extrapolated to other species such as mammals, we focused on mouse embryogenesis in this study. Basically, we obtained similar results to those with *Xenopus* and found that cytoplasmic condensates of ZO-1 exist in the trophectoderm (TE) cells of early mouse embryos before E4.0, although the condensate disappears upon stretching of TE cells due to the expansion of blastocyst cavity (blastocoel).

To confirm that ZO-1 condensates are the product of cellular LLPS (Brangwynne, 2013), we used cultured cells in which GFP-ZO-1 condensates were observed in various conditions. Following both the treatment of a cell-permeabilizing agent digitonin (Shiina, 2019) and analyses with fluorescent recovery after photobleaching (FRAP), ZO-1 condensates showed to have typical properties of liquid droplets generated by LLPS. We also identified an intrinsically disordered region (IDR) (Kato et al., 2012; Li et al., 2012), an unstructured stretch of an amino acid sequence of low complexity that resides in the C-terminal half, and showed that it is necessary for phase separation of ZO-1. Finally, we highlighted the importance of cytoskeletal actin in the regulation of ZO-1 assembly and proposed that ZO-1 protein became liberated from droplets and was deployed to enhance the tight junction in a force-dependent manner. This was supported by observations of MDCK cells in different culture conditions, including wound healing and substrates of different stiffness.

The present study, using early mouse embryos and cultured cells, reveals a previously unknown role of physical force that regulates the condensation of ZO-1, which contributes to the enhancement of cell-to-cell adhesion during early development.

## Results

### ZO-1 condensates in early mouse embryos

In our previous report (Kinoshita et al., 2020), we demonstrated that the stretch of ectodermal tissue in the *Xenopus* gastrula embryo induces a MET-like cellular response, including a reduction of cytoplasmic ZO-1 puncta and its accumulation at the cell junction. To examine whether the behavior of ZO-1 protein is conserved across species, especially in mammals, we first immunostained mouse embryo ZO-1. We focused on E3.5 and E4.5 embryos since they hatch out of the zona pellucida (ZP) and expand their shape (Figure 1A). We fixed E3.5 and E4.5 embryos with paraformaldehyde and immunostained them with a ZO-1 antibody (Figure 1B). We found that the cells of E3.5 embryos showed a significantly higher number of ZO-1 puncta in the cytoplasm relative to E4.5 embryos. As development proceeded, the surface area of TE cells, particularly on the mural side, expanded and became thinner (data not shown). We found that the number of cytoplasmic ZO-1 puncta were reduced, and ZO-1 signal intensity at the plasma membrane in E4.5 embryos became much higher than those of E3.5 embryos at the expense of their cytoplasmic pool (Figure 1C, D), without changing the total fluorescence intensity of the protein (Figure 1E left). Importantly, this change coincides well with the accumulation of F-actin at the cell cortex in E4.5 embryos (Figure 1B and 1E right). This result suggests that the shuttling of ZO-1 protein from the cytoplasmic puncta to cell junctions occurs as development progressed. We observed a similar behavior of ZO-1 protein using ZO-1-EGFP-expressing mouse embryos by live-imaging (Katsunuma et al., 2016) (Figure 1F, G and Video S1). These results are consistent with our previous finding, where ZO-1 puncta decreased in *Xenopus* ectodermal cells that were exposed to a higher tensile force (Hashimoto et al., 2019; Kinoshita et al., 2020).

**Figure 1.**
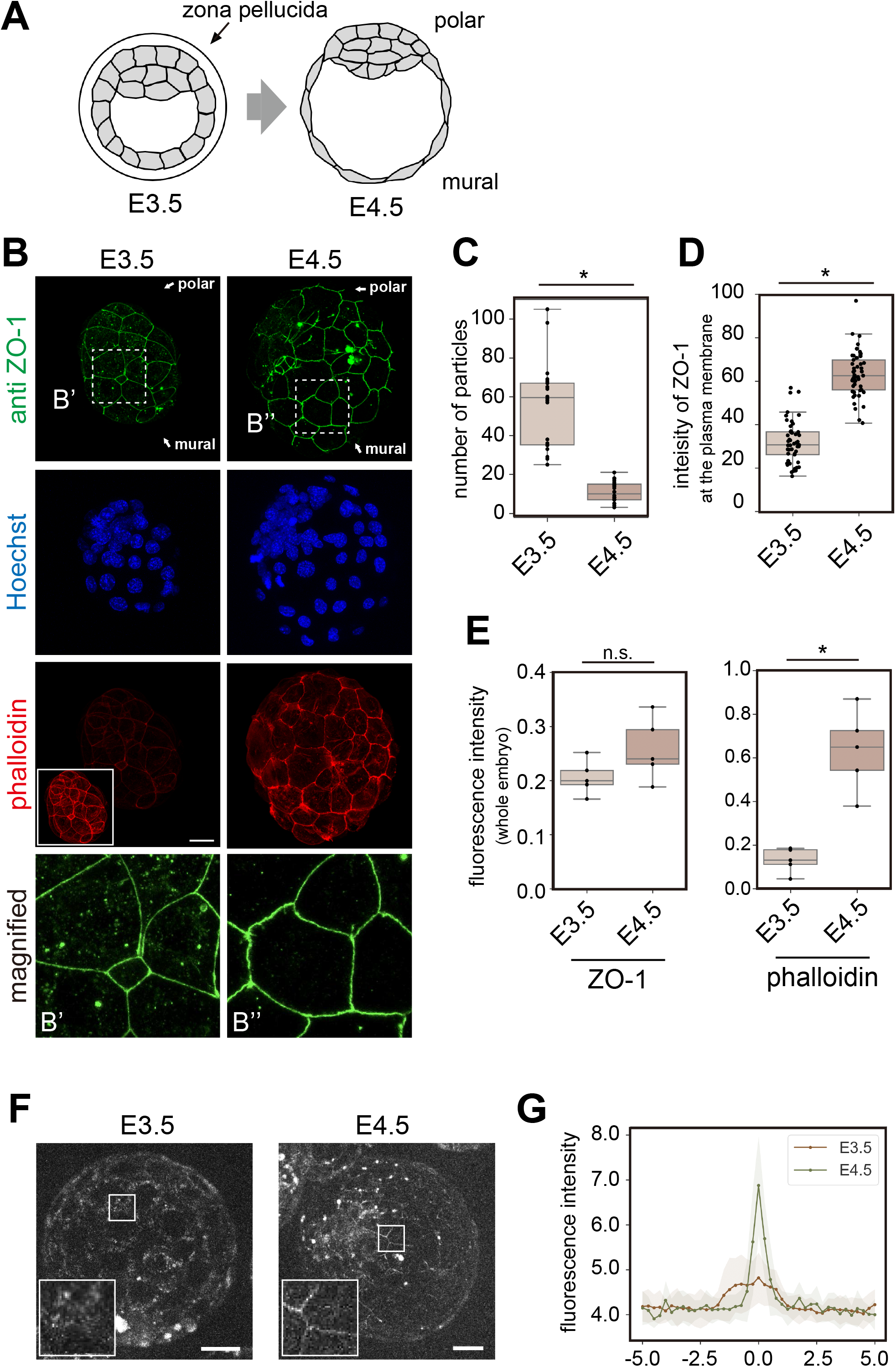
Change of ZO-1 localization in mouse hatching embryos. **A**. Schematic diagram of the hatching process of the mouse embryo. At E3.5, the embryo is covered with the zona pellucida (ZP). During hatching, the embryo is enlarged with an expansion of the trophectoderm (TE), and emerges from the ZP. **B**. Immunofluorescence of mouse E3.5 and E4.5 embryos. Embryos were stained with an anti-ZO-1 antibody, Hoechst 33342, and Alexa Fluor 546 phalloidin. Scale bar, 20 μm. The inset in the E3.5 phalloidin image was acquired with higher laser power, demonstrating that the structure of cortical F-actin is formed at E3.5 even though the signal intensity was weaker than that in the E4.5 embryo. The bottom panels are enlarged images from the upper panels indicated by the dashed squares B’ and B’’. **C**. The number of particles in one TE cell was counted. **D**. The signal intensities of ZO-1 at the cell periphery of the TE at E3.5 and E4.5 were quantified. **E**. Fluorescence intensity of whole embryos stained with the anti-ZO-1 antibody and phalloidin was measured. For comparison, intensities were normalized by Hoechst staining intensity. n = 5 embryos. n.s.: not significant. *p < 0.01 (*t*-test). **F**. Embryos from mice expressing ZO-1-EGFP by EGFP-knock-in were observed. Before hatching (E3.5), ZO-1 puncta were observed around the cell periphery, but cell-cell boundaries were unclear. After hatching (E4.5), membrane localization of ZO-1-GFP became evident. **G**. Fluorescence intensity around cell boundaries in ZO-1-EGFP–expressing embryos were quantified. After hatching, ZO-1-EGFP localized to the plasma membrane and formed sharp boundaries with ZO-1-EGFP. Scale bars, 20 μm.

Interestingly, however, in the polar TE cells attached to the inner cell mass (ICM) of E4.5 embryos, a significant number of condensates still remained in the cytoplasm (Figure S1A and B). Consistently, the fluorescent intensity of both anti-ZO-1 immunostaining and F-actin accumulation at the plasma membrane in polar TE cells was lower than in mural cells (Figure S1C). Since polar TE cells attach to the crowded ICM, these cells might be exposed to lower tensions.

### ZO-1 condensate is regulated by a force-dependent mechanism

We next examined whether the increased tensile force reduced ZO-1 puncta and accumulated it to the junction during hatching. It is known that developing mouse embryos from E4.0 to E4.5 experience a gradually increasing magnitude of luminal pressure due to expansion of the blastocyst cavity in the presence of the ZP (Chan et al., 2019; Leonavicius et al., 2018). Accordingly, the cortical tension of TE increases from E3.5 to E4.5 (Chan et al., 2019). To confirm that the disappearance of ZO-1 condensates from the cytoplasm is dependent on a mechanical force (inner hydraulic pressure of blastocyst cavity), using ouabain, we inhibited Na^+^/K^+^ ATPase, which promotes the influx of water and therefore increases the volume of the blastocyst cavity (Figure 2A and B). In these ouabain-treated embryos, a significant number of ZO-1 condensates was retained in the cytoplasm of mural cells of E4.5 embryos (Figures 2C and D), demonstrating that the cells released from this tension failed to trigger the dissolution of ZO-1 condensates.

**Figure 2.**
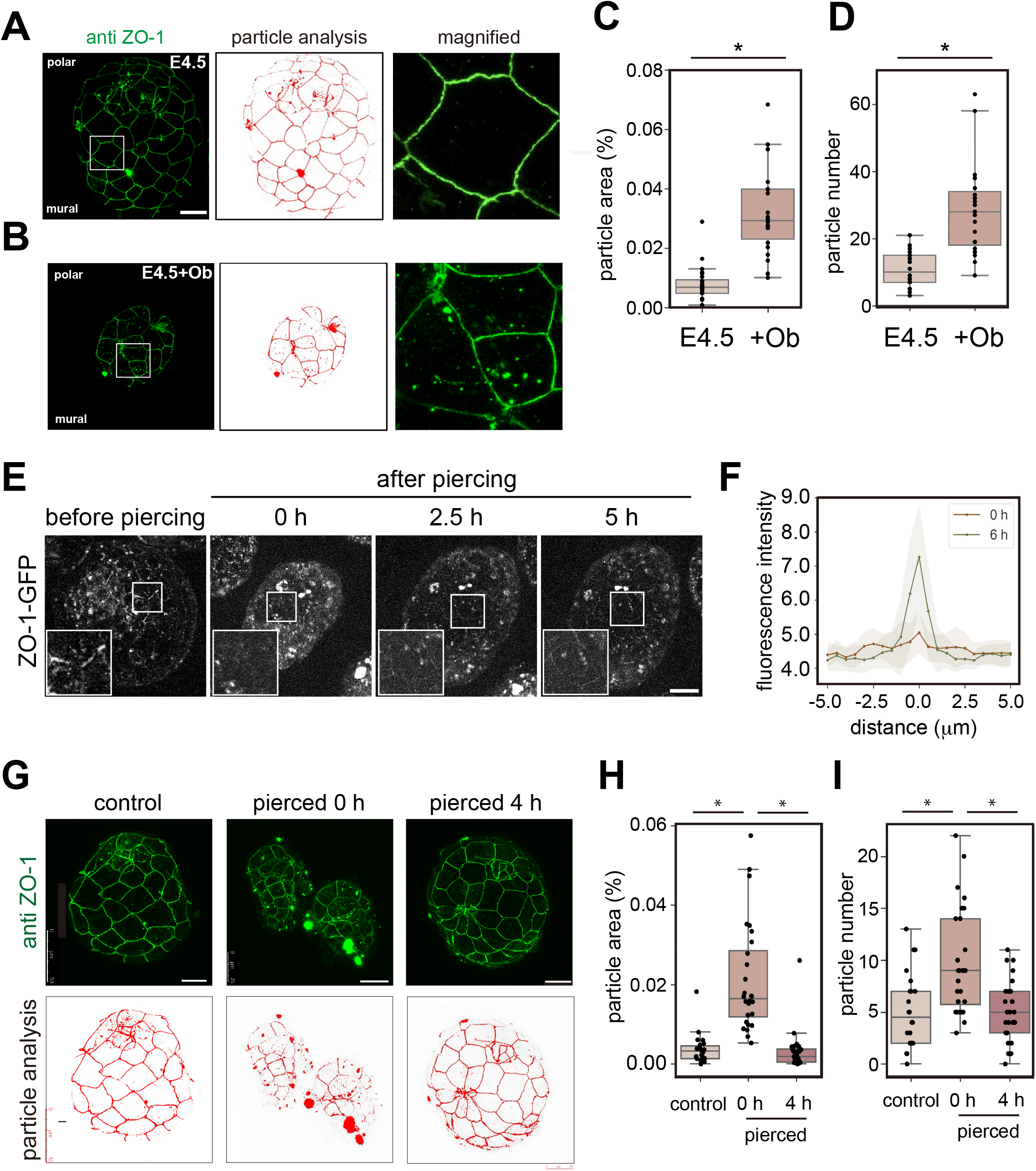
Expansion of TE regulates ZO-1 localization. **A and B**. Embryos were obtained at E3.5 and cultured for 24 h in the absence (A) or presence of 0.3 mM ouabain (+Ob) (B). Embryos were immunostained with the anti-ZO-1 antibody (left), and the images were analyzed using particle analysis in Image J (right). **C** and **D**. The percentage of the particle area (C) and the number of particles (D) of E4.5 mural TE cells were quantified. **E**. E4.5 mouse embryos expressing ZO-1-GFP were pierced with a glass needle. Pierced embryos shrank immediately, and the recovery process was imaged. **F**. The fluorescence intensity of GFP-ZO-1 at the cell periphery at 0 h and 6 h were quantified. The X-axis indicates distance from the cell membrane. *p < 0.01 (*t*-test). **G**. Mouse E4.5 embryos expressing ZO-1-GFP were pierced with a glass needle, then the volume of the blastocyst cavity was reduced, and embryos shrank. Those embryos were fixed and stained with the anti-ZO-1 antibody and fluorescent phalloidin. **H**. The area of particles and cytoplasm of one mural TE cell were measured by Image J software, and the ratio of those values was plotted. **I**. The number of particles was counted using Image J particle analysis. *p < 0.01 (*t*-test). Scale bars, 20 μm.

Next, we mechanically reduced tension by releasing the blastocoel fluid of the ZO-1-EGFP-expressing embryos by piercing embryos with a glass needle. Immediately after they were pierced, embryos shrank, and membrane localization of ZO-1 was reduced, although it recovered within several hours after the wound site had healed and the embryo regained its original shape (Figure 2E, F and Video S2). Immunostaining with the anti-ZO-1 antibody confirmed that endogenous ZO-1 also behaves similarly, observing more cytoplasmic puncta in pierced E4.5 embryos than in control E4.5 embryos (Figure 2G-I). Together, these results suggest that the inner pressure of the growing blastocyst cavity increases the tensile force on TE cells in E4.5 embryos, thereby reducing the number and volume of ZO-1 puncta in the cytoplasm.

This dynamic behavior of ZO-1 protein led us to test whether the ZO-1 puncta in E3.5 embryos were formed by phase separation, treating embryos with 1,6-hexanediol, which is known to dissolve the LLPS assembly (Kroschwald et al., 2015; Shulga and Goldfarb, 2003). As shown in Figures 3A and B, 5% 1,6-hexanediol reduced the number of particles within a few minutes, suggesting that phase separation forms these droplets in the mouse embryos.

**Figure 3.**
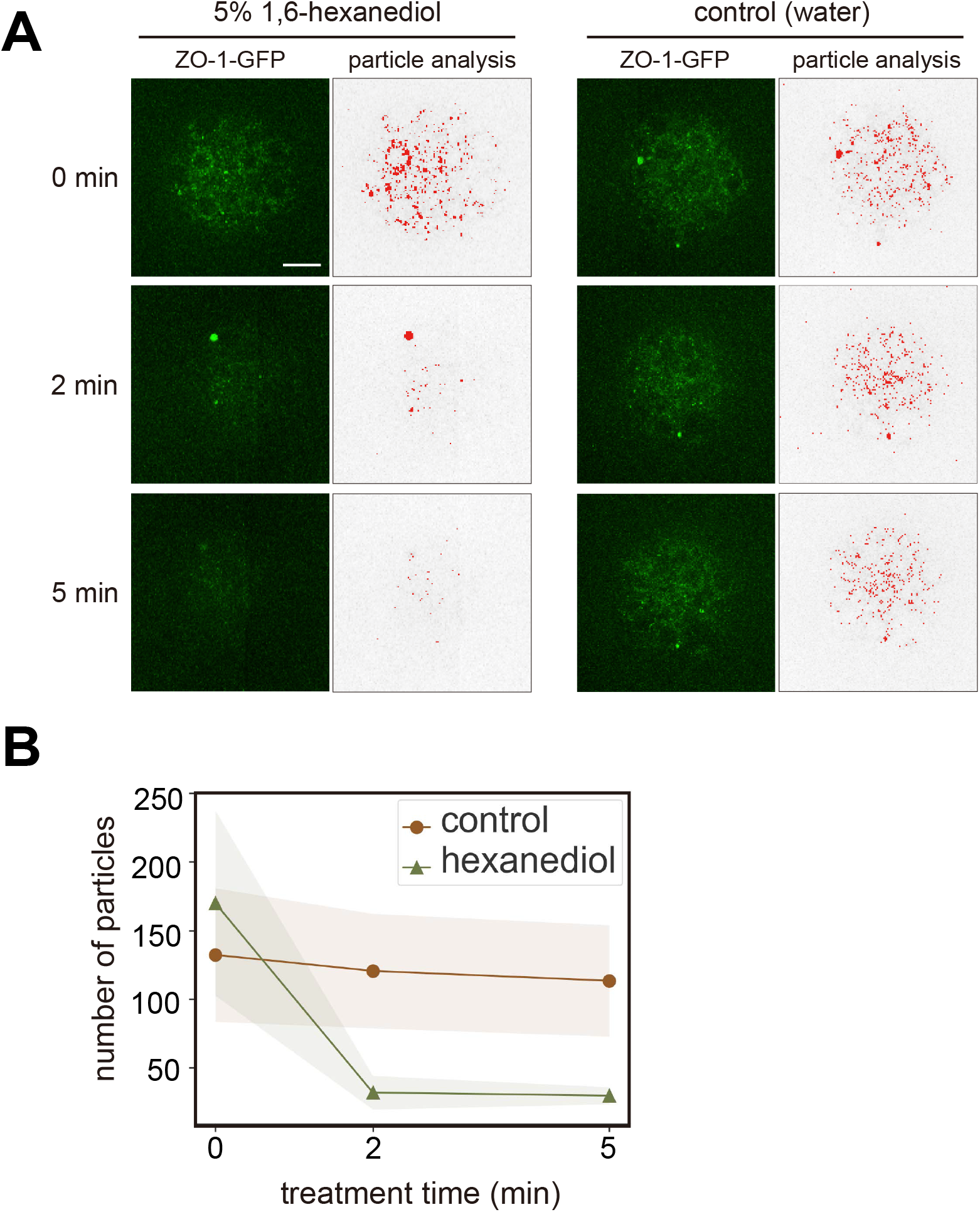
1,6-Hexanediol treatment dissolved ZO-1 droplets in the mouse E3.5 embryo. **A**. Mouse E3.5 embryos were treated with 5 % 1,6-hexanediol. The images were analyzed by particle analysis in Image J (right panels). **B**. The number of particles was counted. n = 5 (hexanediol) and 7 (control) embryos.

### LLPS forms ZO-1 droplets in MDCK cells

In order to analyze the nature of the ZO-1 condensate in various cell environments, we performed a series of studies using canine MDCK cells. The results using mouse embryos suggested that tensile force to the TE cells negatively regulates ZO-1 phase separation. In MDCK cells, it had been demonstrated that increased substrate stiffness increases tension acting on ZO-1 (Haas et al., 2020). We thus examined whether substrate stiffness affects ZO-1 droplet formation. We plated MDCK cells on a polyacrylamide gel substrate of different stiffnesses and immunostained ZO-1. As shown in Figures 4A and B, cells on the soft substrates formed more cytoplasmic ZO-1 puncta than cells on the rigid substrates. This result supports our notion that sensing mechanical stimuli, including through focal adhesion, triggers the remodeling of ZO-1.

**Figure 4.**
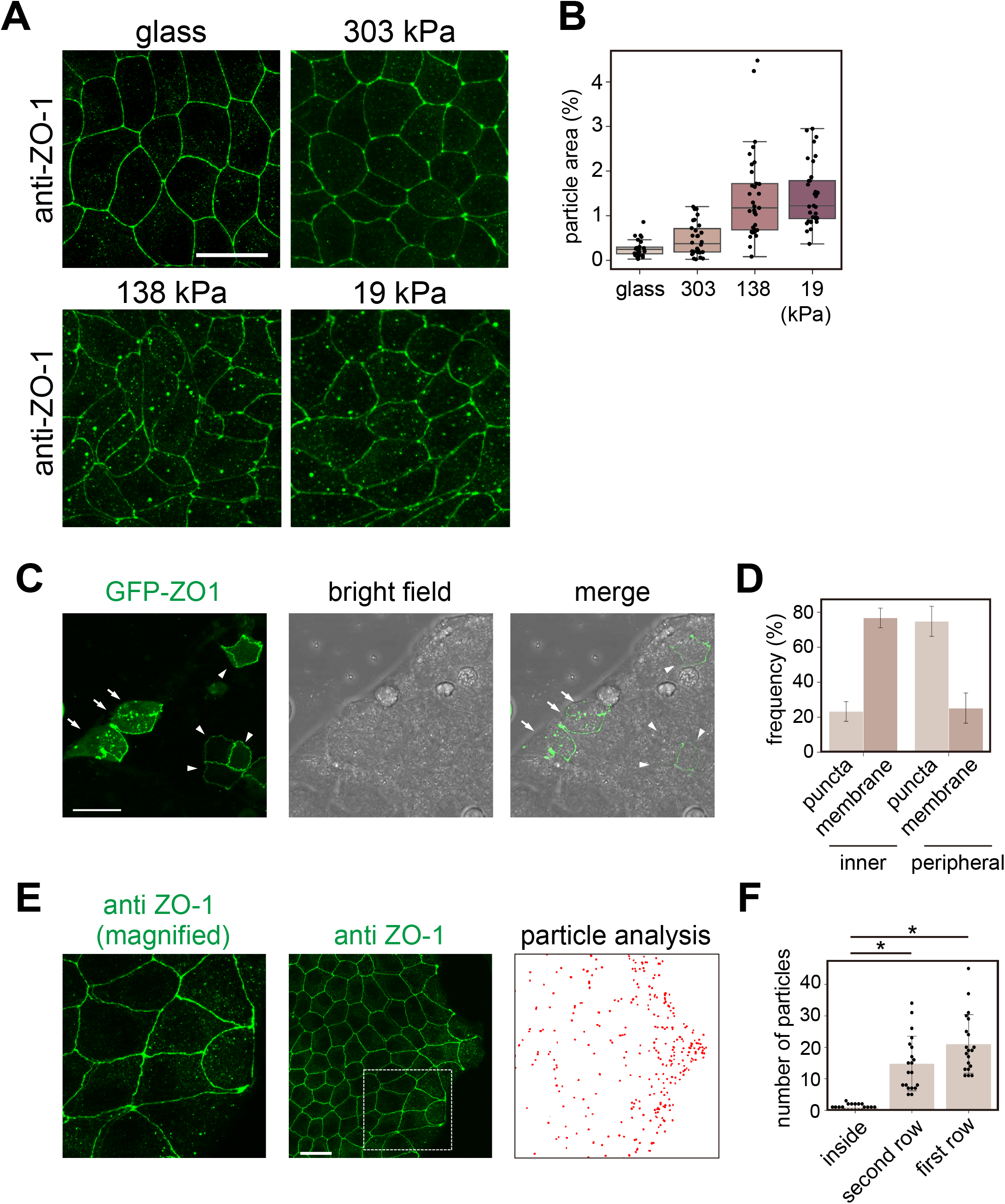
1,6-Hexanediol treatment dissolved ZO-1 droplets in the mouse E3.5 embryo. **A**. MDCK cells were grown on glass or on a polyacrylamide gel with 303, 138, and 19 kPa elasticity. **B**. The ratio of the particle area in each cell was counted by Particle Analysis in Image J. **C** and **D**. GFP-ZO-1 was expressed in MDCK cells, forming islands. In cells located inside the islands (indicated by arrowheads), GFP-ZO-1 was localized mainly at the plasma membrane. In contrast, in cells at the periphery or outside of the colony (indicated by arrows), GFP-ZO-1 formed puncta. **E**. MDCK cells that formed a colony were immunostained with the anti-ZO-1 antibody. Images were analyzed using Particle Analysis in Image J. **F**. Cells in the island were divided into three groups with respect to their location, namely the first and second rows from the edge and the central region of the island. The number of particles in each cell was counted. n = 20 cells. *p < 0.01 (*t*-test).

We also queried whether ZO-1 localization would change in an interaction with surrounding cells, which might affect the mechanical environment (Zhu et al., 2000). We grew MDCK cells to form islands in a culture dish and transfected a plasmid harboring human ZO-1 fused to GFP (GFP-ZO-1). It is notable that cells in the central region lost condensates while peripheral cells tended to have condensates (Figure 4C and D). However, we occasionally observed cells that harbored relatively large condensates in the central region, but those cells were not well attached to, and were free from, neighboring cells (data not shown). This tendency was confirmed by immunofluorescence using the anti-ZO-1 antibody (Figure 4E). Quantitatively, cells in the forefront row (most outer layer) and the second row had 15 to 20 condensates per cell on average. In contrast, cells in the inner or central region had only a few condensates (Figure 4F). One feature that distinguishes the inner cells from peripheral cells is that after cells reach confluency, the former adopts polygonal shapes and has higher densities than the latter. These results suggest that the inner cells establish robust cell-to-cell contact by tight and adherens junctions, which are physically supported by the actin cytoskeleton, and an individual cell rigidly contacts with surrounding cells, resembling embryonic cells under a tensile force.

To confirm further that the ZO-1 droplets observed in MDCK cells are indeed the products of LLPS, we treated cells harboring GFP-ZO-1 droplets with digitonin, a cell-permeabilizing reagent that disrupts the equilibrium between a condensate and the cytoplasm. That treatment shrank the droplets within several minutes in MDCK cells (Figure 5A, B and Video S3). Furthermore, we performed FRAP analyses on the GFP-ZO-1 droplets in MDCK cells (Figure 5C and D). The fluorescent signal of GFP-ZO-1 droplets recovered by almost 50% within 2 min. These results demonstrate that the ZO-1 puncta are droplet-like condensates produced by LLPS and are in equilibrium with the cytoplasm.

**Figure 5.**
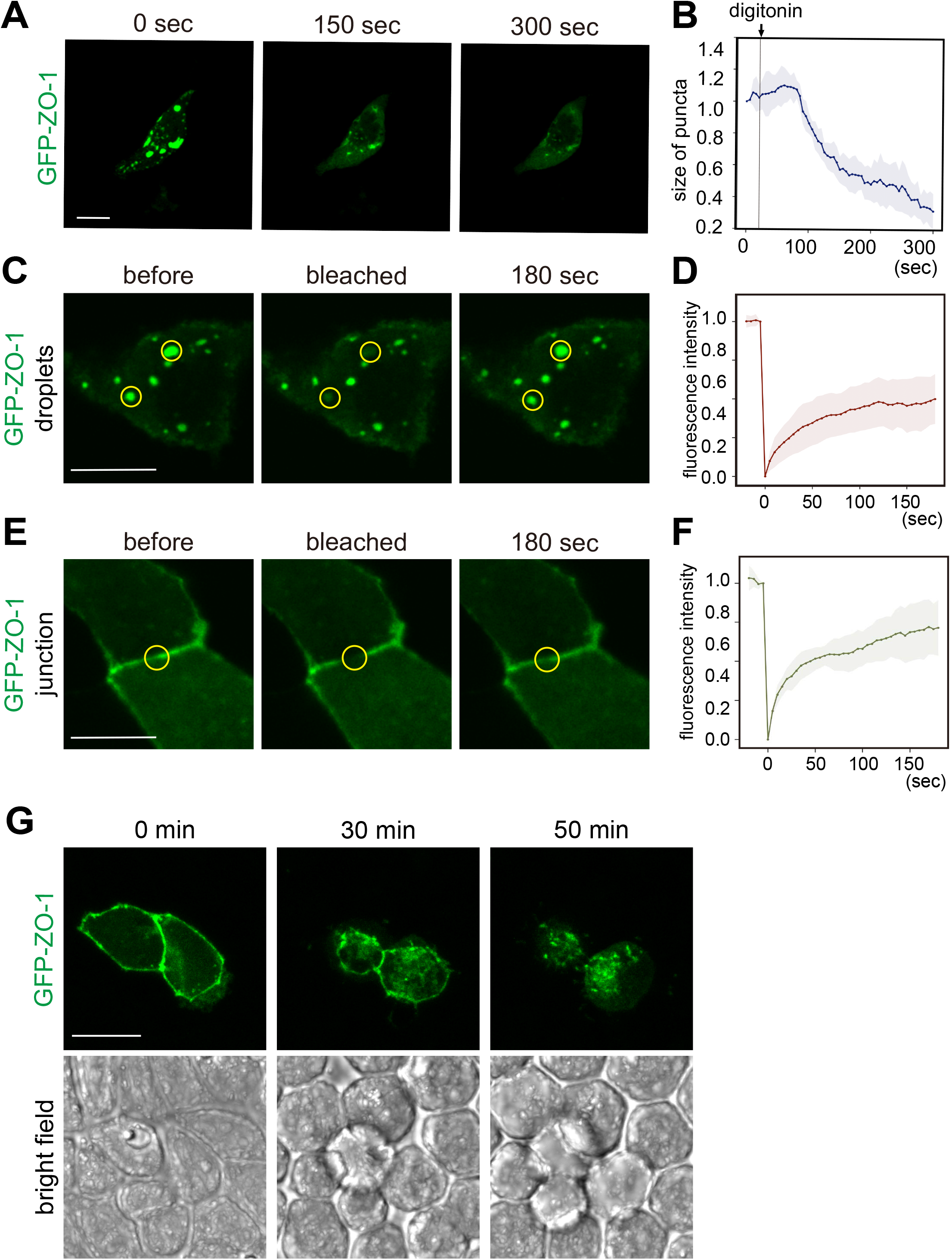
Dynamics of ZO-1 puncta in MDCK cells. **A** and **B.** MDCK cells expressing GFP-ZO-1 were semi-permeabilized with 0.012% digitonin. The size of puncta was quantified with Image J. n = 12 puncta. **C** - **F**. Fluorescent recovery after a photobleaching (FRAP) assay. **C** and **D**. The recovery of fluorescence of puncta was quantified. n = 20 puncta. **E** and **F**. The recovery of junctional ZO-1 was quantified. **G**. MDCK cells were treated with trypsin-EDTA solution. As cells lost their cell-cell contact and their shape became spherical, ZO-1 detached from the plasma membrane and formed puncta. Scale bars, 20 μm.

It has been reported that junctional ZO-1 is also assembled by the LLPS (Beutel et al., 2019). We thus conducted a FRAP assay using inner cells with junctional ZO-1 in the same culture dish (Figure 5E and F) and found that ZO-1 at the junction showed a similar recovery rate. These results indicate that both ZO-1 in the cytoplasmic droplets and ZO-1 at the junction are liquid-like condensates and that ZO-1 changes its localization depending on the cellular environment.

### Cell-cell interaction regulates ZO-1 droplet formation

Our finding that peripheral cells have more droplets than inner cells suggested that robust cell-cell contact negatively regulates ZO-1 LLPS. To verify this idea, we reduced cell-to-cell contact by treating MDCK cells expressing GFP-ZO-1 with PBS containing trypsin-EDTA. As expected, when trypsin-EDTA solution was added, ZO-1 detached from the cell membrane and formed puncta as cells lost their contact (Figure 5G and Video S4).

The robustness of cell-cell contact was also examined by laser ablation. MDCK cells were grown so that they formed islands in one culture dish. After the GFP-ZO-1 expression plasmid was transfected, cells were irradiated with laser light. When cells inside the colonies were irradiated at the cell junction, cells stably stayed after irradiation due to cell-cell contact with neighboring cells (Figure S2A, C and Video S5). When peripherally-located cells with ZO-1 puncta were irradiated, those cells immediately erupted and disappeared after irradiation (Figure S2B, C, and Video S6), indicating that those cells are weakly attached to surrounding cells. These results suggest that in MDCK cells, the formation of ZO-1 condensate is negatively associated with the establishment of robust cell-to-cell contact.

### ZO-1 forms cytoplasmic condensates in migrating cells during wound healing

To further examine whether cell-to-cell contact negatively regulates ZO-1 condensate formation in the cytoplasm of MDCK cells, we conducted a wound-healing assay using GFP-ZO-1–expressing cells (Figure 6A and B and Video S7). Initially, most of the cells had few condensates in a confluent cell sheet. After scratching, cells started active migration toward the wound site, forming a significant number of droplets. By longer incubation in this assay, cells migrating from both sides eventually met in the middle of the gap, re-establishing a new cell-cell interaction. During this process, ZO-1 droplets disappeared and accumulated at the cell junction (Figure S3 and Video S8). These results indicate that ZO-1 condensates shuttle between cytoplasmic droplets and the cell junction, depending on cell-cell interaction.

**Figure 6.**
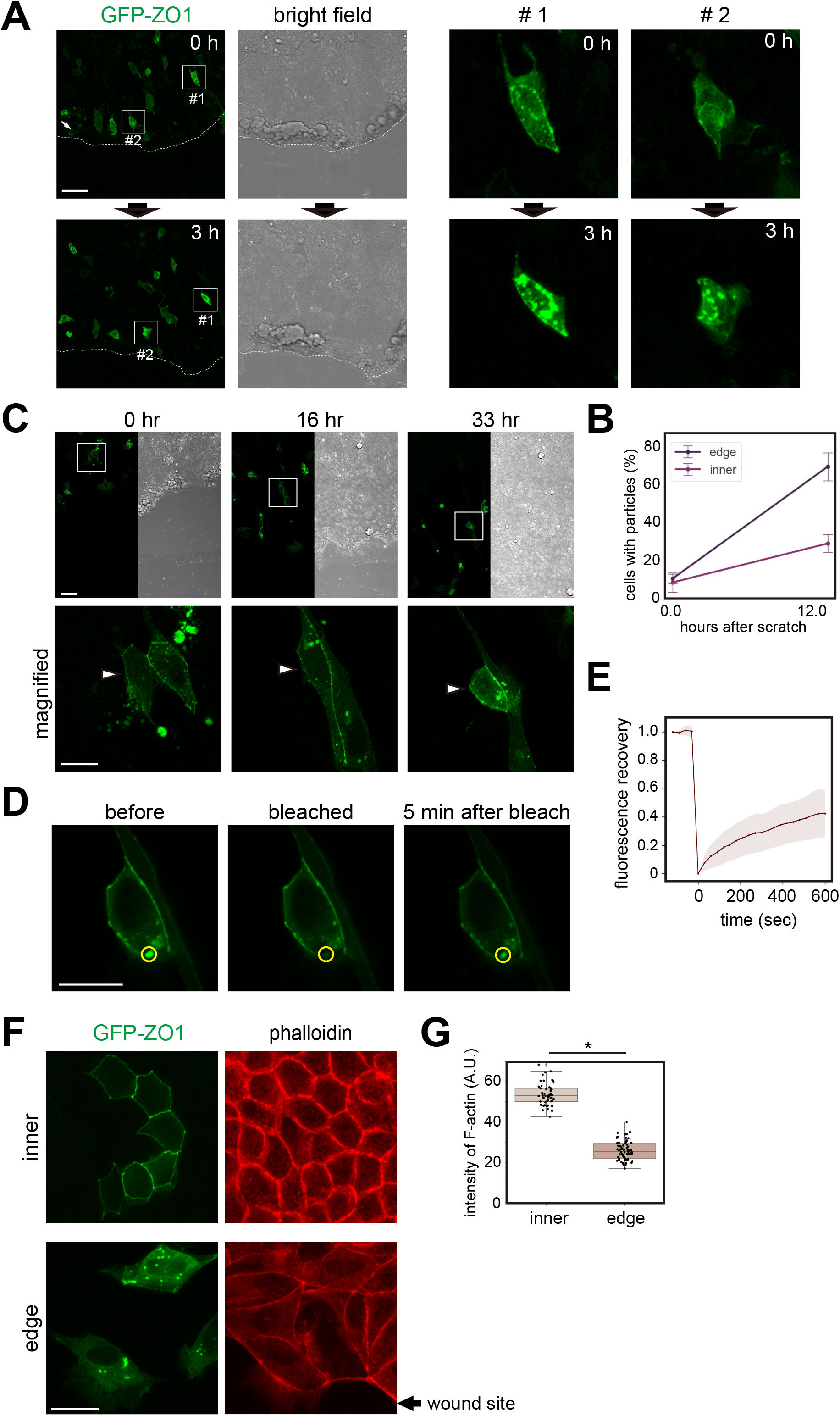
Wound-healing assay using MDCK cells expressing GFP-ZO-1. **A** and **B**. The confluent cell sheet was scratched and observed. Immediately after scratching (0 h), GFP-ZO-1 was at the cell periphery, whereas after 3 hours (3 h), it formed puncta in many of the cells close to the scratch (within 200 μm). The right panels were magnified from #1 and #2 cells in **A**. Scale bar, 50 μm. **B**. The number of cells with GFP-ZO-1 particles was counted. Cells located within 200 μm from the scratch site were defined as “edge” cells. Cells more than 200 μm away from the scratch site were defined as “inner” cells. Data were obtained from three experiments. **C-E**. Wound-healing assay of MDCK cells expressing GFP-ZO-1 followed by a FRAP assay. Cells formed GFP-ZO-1 puncta in the wound-healing assay were subjected to the FRAP assay. Three cells were analyzed. Scale bars, 20 μm. **F**. Inner and edge cells were fixed 10 h after scratching and stained with phalloidin. Arrow indicates the wound site: scale bar, 20 μm. **G**. Fluorescence intensity of phalloidin staining at the plasma membrane was measured. *p < 0.01 (*t*-test).

The FRAP assay revealed that the fluorescent signals of GFP-ZO-1 condensates that had formed during the wound-healing assay recovered to about 40% within 10 min, suggesting that these condensates were formed by LLPS (Figure 6C-E, Video S9, and S10). This result again indicates that the formation of ZO-1 condensate is regulated by intercellular and extracellular environments such as cell-to-cell contact.

In the wound-healing assay, we found that the distribution and intensity of F-actin were different between inner cells and edge cells. As shown in Figures 6F and G, cortical F-actin was well-developed in the non-migrating polygonal cells with junctional ZO-1 (“inner” cells in Figure 6F). In contrast, migrating cells around the wound site, which form ZO-1 droplets, had reduced cortical actin and developed more stress fibers. This correlation between ZO-1 and F-actin is consistent with our previous finding in which the application of force to *Xenopus* embryonic cells induced a MET-like response, including the accumulation of cortical F-actin, and reduced ZO-1 condensates as cell-cell junctions were enhanced (Hashimoto et al., 2019; Kinoshita et al., 2020).

### Interaction with F-actin regulates ZO-1 phase separation

We further investigated the relationship between ZO-1 phase separation and F-actin organization by expressing GFP-ZO-1 in non-confluent *Xenopus* A6 cells. In A6 cells, GFP-ZO-1 localized mostly in the cell periphery and colocalized with F-actin bundles (Figure 7A). After treating the cells with latrunculin B, a potent inhibitor of actin polymerization, ZO-1 granules rapidly emerged in the cytoplasm, which disappeared soon after washing out the drug (Figure 7B and Video S11). These results suggest that the ZO-1 granules, which were highly mobile in the FRAP assay (Figure 7C and D), are negatively regulated by the interaction with F-actin in A6 cells. We also treated MDCK cells expressing GFP-ZO-1 with latrunculin B, but no detectable change in the localization of ZO-1 was observed (data not shown). Therefore, we speculate that the contribution of F-actin to suppress cytoplasmic ZO-1 condensation is more significant in A6 cells than in MDCK cells, which develop more rigid junctional structures.

**Figure 7.**
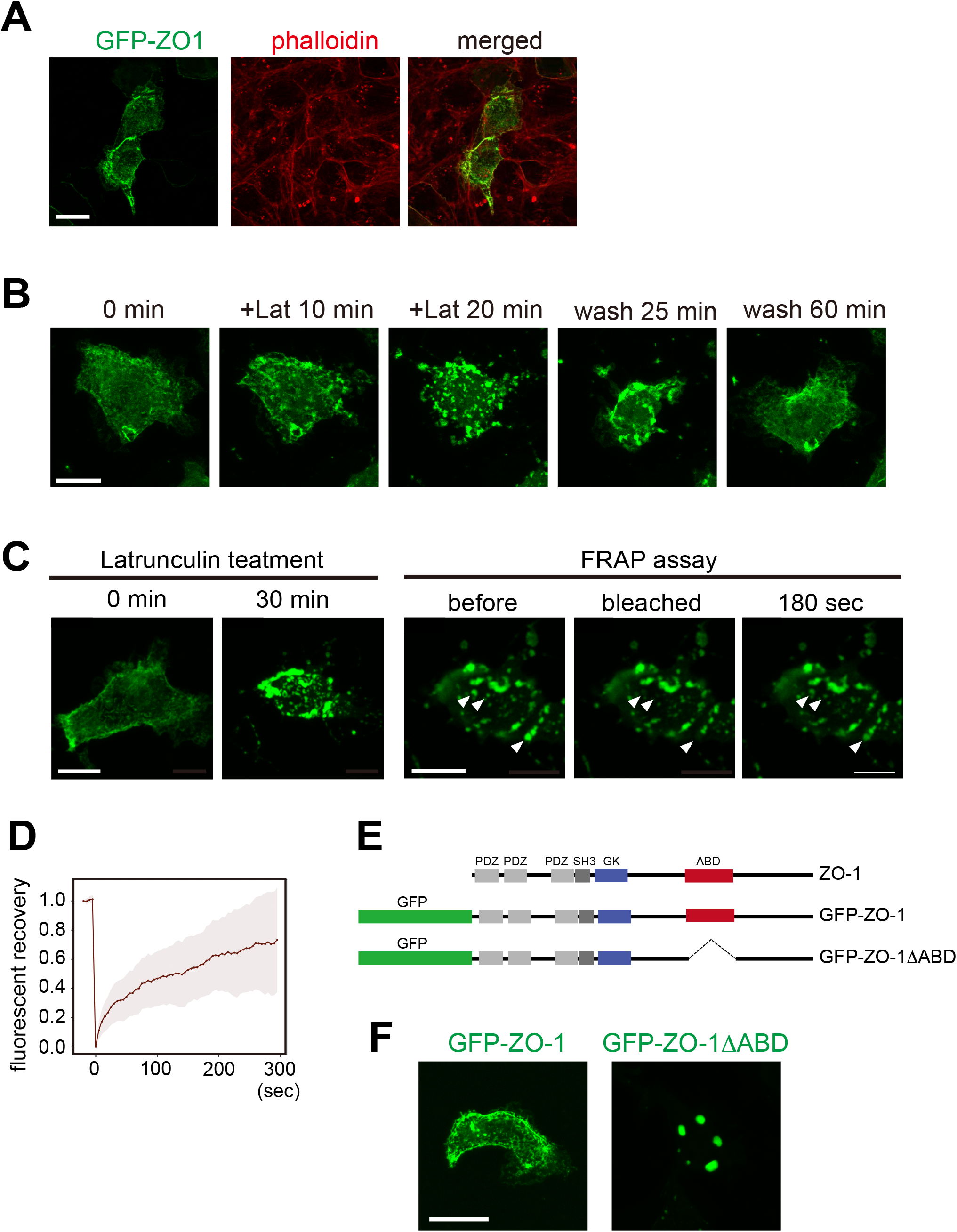
GFP-ZO-1 binding to F-actin in *Xenopus* A6 cells. **A**. GFP ZO-1 was transiently expressed in *Xenopus laevis* A6 kidney cells. Cells were fixed and stained with fluorescent phalloidin. **B**. A6 cells expressing GFP-ZO-1 were treated with 0.5 μM latrunculin B for 20 min, then washed out. **C**. A6 cells were treated with 0.5 μM latrunculin B for 30 min. Then, a FRAP assay was performed. Granules, indicated by arrowheads, were bleached. **D**. The recovery rate was quantified (n = 14). **E**. Domain structure of ZO-1 protein and expression constructs used here: GFP-tagged full-length human ZO-1 (GFP-ZO-1), actin-binding domain (ABD)-deletion mutant (GFP-ZO-1ΔABD). **F**. Localization of GFP-ZO-1 and GFP-ZO-1ΔABD. Scale bars, 20 μm.

To understand the F-actin-driven mechanism, we constructed GFP-ZO-1ΔABD (Figure 7E), which lacks the actin-binding domain (ABD) (Fanning et al., 2002), and expressed it in A6 cells. As shown in Figure 7F, GFP-ZO-1ΔABD formed droplets with a smooth surface. This suggests that cytoplasmic droplet formation is restricted by the binding of ZO-1 to F-actin (Beutel et al., 2019; Schwayer et al., 2019) and that GFP-ZO-1ΔABD lost its capacity for the interaction. Supporting this notion, co-expression with a red fluorescent protein fused to F-actin-binding peptides, Lifeact, Utrophin, and the actin-binding domain of Moesin (MoesinABD), similarly induced condensates that were indistinguishable from those induced by GFP-ZO-1ΔABD (Figure S3A). It is likely that the expression of these actin-binding proteins competes binding to F-actin with ZO-1.

We also observed that the MoesinABD-induced ZO-1 condensates moved dynamically, sometimes fused and underwent fission (Figure S4B and Video S12), reminiscent of typical liquid droplets. In addition, the size of these droplets was rapidly reduced by cell permeabilization following digitonin treatment (Figure S4C and Video S13) and exhibited a rapid recovery of fluorescence in the FRAP assay (Figure S4D). These results confirm that these cytoplasmic droplets are formed by phase separation.

### The C-terminal intrinsically disordered region is required for ZO-1 phase separation

It is known that intrinsically disordered regions (IDRs) are required for proteins to undergo phase separation (Kato et al., 2012; Li et al., 2012). Four IDRs are predicted in human ZO-1 (Figure S5A) using the Protein DisOrder prediction System (https://prdos.hgc.jp/cgi-bin/top.cgi) (Ishida and Kinoshita, 2007). In order to address which IDR is critical for ZO-1 in A6 cells, we expressed a series of deletion mutants (Figure S5B). As a positive control, we expressed GFP-ZO1-ΔABD and confirmed that it formed droplets. Then, we deleted IDR1, IDR2, and IDR3, which reside in the N-terminal half, from GFP-ZO1-ΔABD (GFP-ZO1-ΔABD-ΔIDR123). When it was expressed, it also formed droplets (Figure S5C), indicating that these regions are not required for phase separation in A6 cells. In contrast, deletion of the C-terminal IDR4 (GFP-ZO1-ΔABD-ΔIDR4) abolished ZO-1 droplets, and it became dispersed in the cytoplasm. The result demonstrates that IDR4 is critical for cytoplasmic liquid droplet formation by LLPS. Since the actin-binding domain overlaps with IDR4, it is possible that F-actin binding may affect the LLPS-inducing activity of IDR4.

## Discussion

Our present study using mouse embryos as well as cultured cells, together with a previous study that employed *Xenopus* embryos (Kinoshita et al., 2020), indicates that similar mechanisms may govern cellular responses to mechanical forces across species. Briefly, mechanical stresses caused by dynamic morphogenesis and/or vigorous cell movements during development, such as gastrulation, activate the force-dependent pathway leading to the remodeling of cell-to-cell junctions. The present study also suggests that the process involves the negative regulation of cytoplasmic ZO-1 condensate formation by LLPS, leading to its translocation to the cytoplasm. Since embryonic tissues, as well as some adult tissues are constantly exposed to various forces, and since ZO-1 is a structurally conserved protein among species, we propose that this mechanism may be employed to maintain the robustness and integrity of tissues against various forces in general, and might also have been an evolutionary innovation when multicellular animal species such as placozoa emerged (González-Mariscal et al., 2011). Although our previous study using *Xenopus* embryos identified that the function of the FGF receptor is essential for the force-induced pathway, it also suggested that FGF ligands are not required for signal activation, shedding light onto the as yet unrevealed activation mechanisms for FGFRs and other receptor tyrosine kinases (RTKs) by force. In fact, the epidermal growth factor receptor (EGFR), which is an upstream RTK for Erk, was implicated in the mechanical force-dependent activation of Erk in the collective migration of wound-healing cells (Hino et al., 2020). In both cases, cytoskeletal remodeling by tensile forces appears to trigger these pathways, supporting the idea that mechanoresponsive signaling pathways may be highly conserved.

Interestingly, our present study using cultured cells demonstrated that the efficacy of ZO-1 condensation as droplets largely depends on cell type. In MDCK cells, cytoplasmic ZO-1 condensates are formed when they are cultured sparsely, or cell-cell interaction is weakened, for example, at the periphery of the colonies, during the wound healing process, or by the trypsin/EDTA treatment. In contrast, in cells with a robust cell-cell interaction, ZO-1 condensates appear to be localized at the cell junction. In the case of A6 cells, blocking the interaction between ZO-1 and F-actin induces ZO-1 droplet formation, suggesting that ZO-1 droplet formation is negatively regulated by the interaction with F-actin. This suggests that the difference in the amount or dynamics of F-actin may distinguish these two cell types and the revelation of ZO-1 from F-actin is critical for maintaining large-sized ZO-1 condensate.

It is tempting to speculate that Erk might regulate F-actin polymerization through small G proteins such as GEF-H1 and its downstream target RhoA (Fujishiro et al., 2008; Itoh et al., 2014; Ren et al., 1998). Recently, Haas et al. (2020) reported that substrate stiffness regulates GEF-H1, thereby increasing tension, by acting on ZO-1. Since we reported in this study that substrate stiffness changes ZO-1 droplet formation, it would be intriguing to investigate whether the GEF-H1 pathway is involved in the regulation of ZO-1 phase separation. Second, as described above, it was reported that Erk is activated in collectively migrating MDCK cells by EGFR and regulates F-actin organization and cellular contraction (Hino et al., 2020). It is also possible that the cascade of phosphorylation initiated by RTKs might eventually result in the remodeling of F-actin and ZO-1 localization.

It has been reported that the assembly of ZO-1 at the cell junction is also driven by phase separation (Beutel et al., 2019). We also confirmed that both the cell junction and the cytoplasmic ZO-1 granules show typical characteristics of phase-separated condensates, which suggests that ZO-1 localization may be dynamically switched depending on the mechanical environments and F-actin plays a key role in the switching in A6 cells. We also mapped the region essential for cytoplasmic granule formation to the fourth IDR (IDR4 in Figure S4). As IDR4 covers the actin-binding domain, the interaction with F-actin may directly affect its phase separation. It has been reported that F-actin binding to ZO-1 induces its conformational change (Spadaro et al., 2017). Such structural changes may regulate the phase separation of IDR4 to form cytoplasmic droplets. Beutel et al. (2019) demonstrated that the PDZ-SH3-GuK domain in ZO-1 plays an essential role in phase separation at the cell junction. Since ZO-1 is a scaffold protein that interacts with many proteins (Fanning and Anderson, 2009), such a protein-protein interaction may also change the state of phase separation, resulting in the difference between the cytoplasmic and junctional forms of condensates. Investigating the precise molecular mechanisms that could explain the two distinct states of ZO-1 will be important.

It has been reported that tension mediates the remodeling of cell junctions (Ito et al., 2017). Phase separation of ZO-1 may be directly involved in this regulatory system. In this scenario, LLPS may act as an on-demand ON/OFF switch of ZO-1 supply to the cell membrane from the cytoplasm, and this mechanism may be shared and employed by multicellular animal species. Therefore, its evolutionary origin is an intriguing problem worth studying in the future.

ZO-1 is implicated in various pathologies, including cancer. Particularly, in EMT, the loss of cell-to-cell adhesion is known to drive the progress and metastasis of cancer. It is also known that in neuronal cells, some condensates which contain proteins, such as the RNA-binding protein FUS (Murray et al., 2017) and tau implicated in Alzheimer’s disease (Wegmann et al., 2018), become aggregated depending possibly on their concentrations, sometime after their formation and/or their chemical modifications. Those proteins are irreversibly aggregated and become dysfunctional, attenuating the neuronal activity of the cells. Collectively considering these observations, we speculate that impairment of the normal regulation of the LLPS of ZO-1 due to changes in intracellular or extracellular conditions could be a possible cause of cancer pathology. Therefore, in addition to embryogenesis, investigating the behavior of ZO-1 in adult tissues, especially in pathological conditions, would deepen our understanding of the physiological significance of the ZO-1 reservoir in cells.

### Limitations of this study

We propose in this paper that F-actin remodeling is one of the key factors to regulate ZO-1 shuttling between cytoplasmic and junctional condensates. However, particularly in MDCK cells, disruption of F-actin did not change junctional ZO-1 localization, suggesting that F-actin remodeling may not be sufficient for re-localizing ZO-1 in MDCK cells. Since reducing cell-cell contact-induced cytoplasmic ZO-1 droplet formation, a mechanism that releases junctional ZO-1 to the cytoplasm must exist. The interactions with their junctional components, such as claudin, occludin, JAM-A, and cingulin (Fanning and Anderson, 2009; Haas et al., 2020; Vasileva et al., 2017) remains to be clarified.

## Acknowledgments

R26-ZO-1-GFP mice were provided by LARGE, BDR (accession # CDB026K), RIKEN, Japan. We thank Dr. Fumio Matsuzaki, RIKEN, for human ZO-1 cDNA, Dr. Makoto Suzuki, Hiroshima University, for the GFP-ZO-1/pCS2 construct, and Dr. Lance Davidson, University of Pittsburgh, for the mCherry-Utrophin construct. We thank Dr. Yuko Mimori-Kiyosue, RIKEN, for A6 cells and Dr. Mitsuru Nishita, Fukushima Medical University, and Dr. Kensaku Mizuno, Tohoku University, for MDCK I cells. We thank the BioImaging Facility, Core Research Facilities of the National Institute for Biology (NIBB), for technical support for live imaging and FRAP analysis. We also thank Dr. Nobuyuki Shiina, NIBB, and Dr. Michael Levine, Princeton University, for technical advice and valuable discussion on LLPS, and insightful comments on this work. This research was supported by JSPS KAKENHI 20K06663 to NK, JSPS KAKENHI 17H03689 and 16H06280 to TF, and MEXT KAKENHI 22127007 and JSPS KAKENHI 15H05865 to NU. TF and NU were also supported by JST CREST JPMJCR1654 and Joint Research of the Exploratory Research Center on Life and Living Systems (ExCELLS), respectively.

## Authors’ Contributions

N.K., T.S.Y., N.Y. and C.T. conducted experiments using tissue culture cells; T.F., N.K., and N.Y. conducted experiments using mouse embryos; N.K. analyzed the data and co-wrote the manuscript. N.U. supervised the research, reviewed the data and conclusions, and wrote the manuscript.

## Supplementary Figure and Video Legends

**Figure S1.**
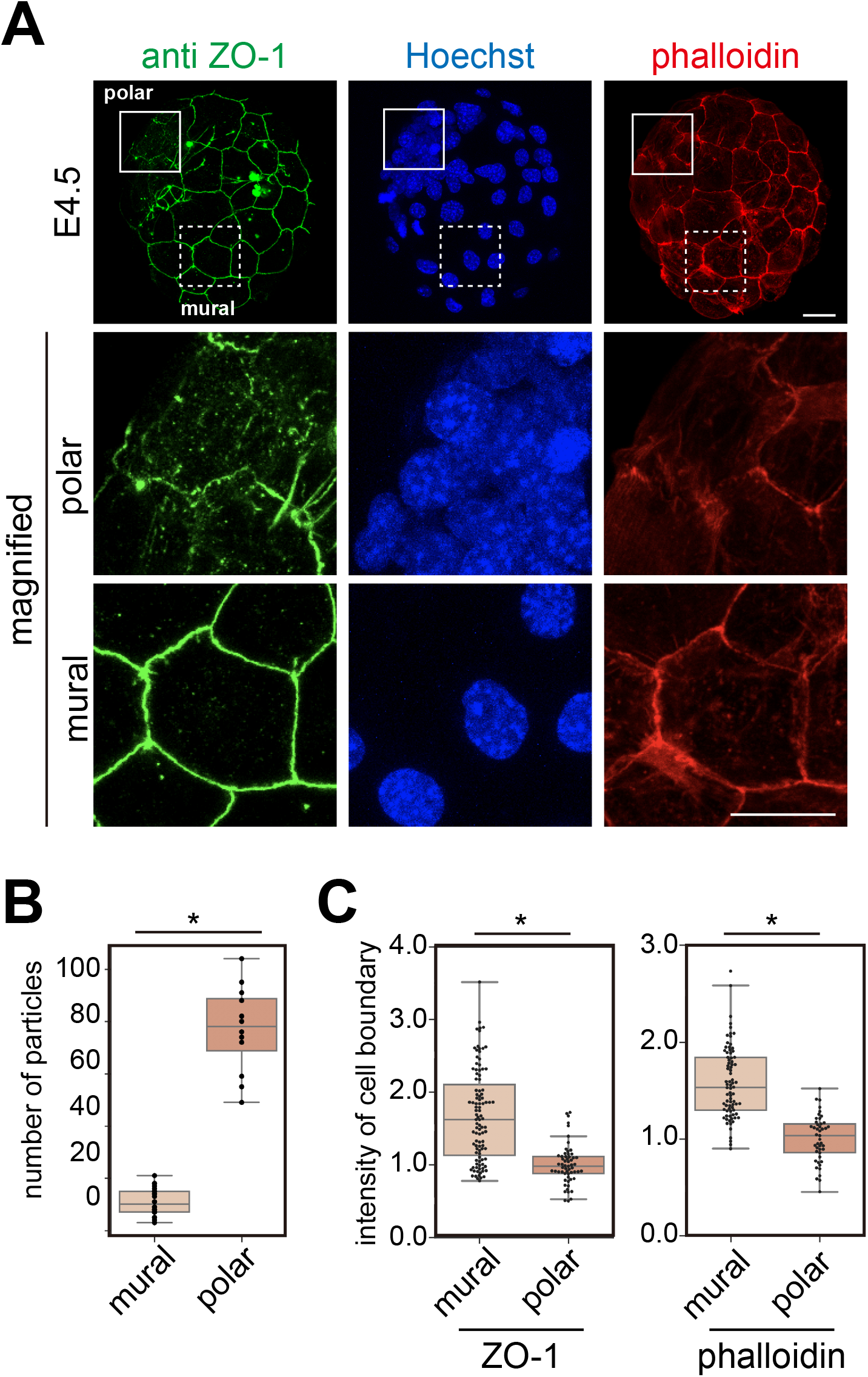
Comparison of ZO-1 and F-actin localization between polar and mural TE cells. **A**. The nuclear-dense inner cell mass (ICM) region was determined by Hoechst33342 staining. ‘Polar’ indicates TE attaching ICM, and ‘mural’ includes TE not attaching the ICM. The upper images are the same as in Figure 1B right images (E4.5). The indicated areas of the middle and the lower images were magnified. Scale bars, 20 μm. **B**. The number of particles in one TE cell was counted. **C**. Signal intensities of ZO-1 and phalloidin at the cell periphery of the mural and polar TE were quantified. 4 - 5 embryos were analyzed for each condition.

**Figure S2.**
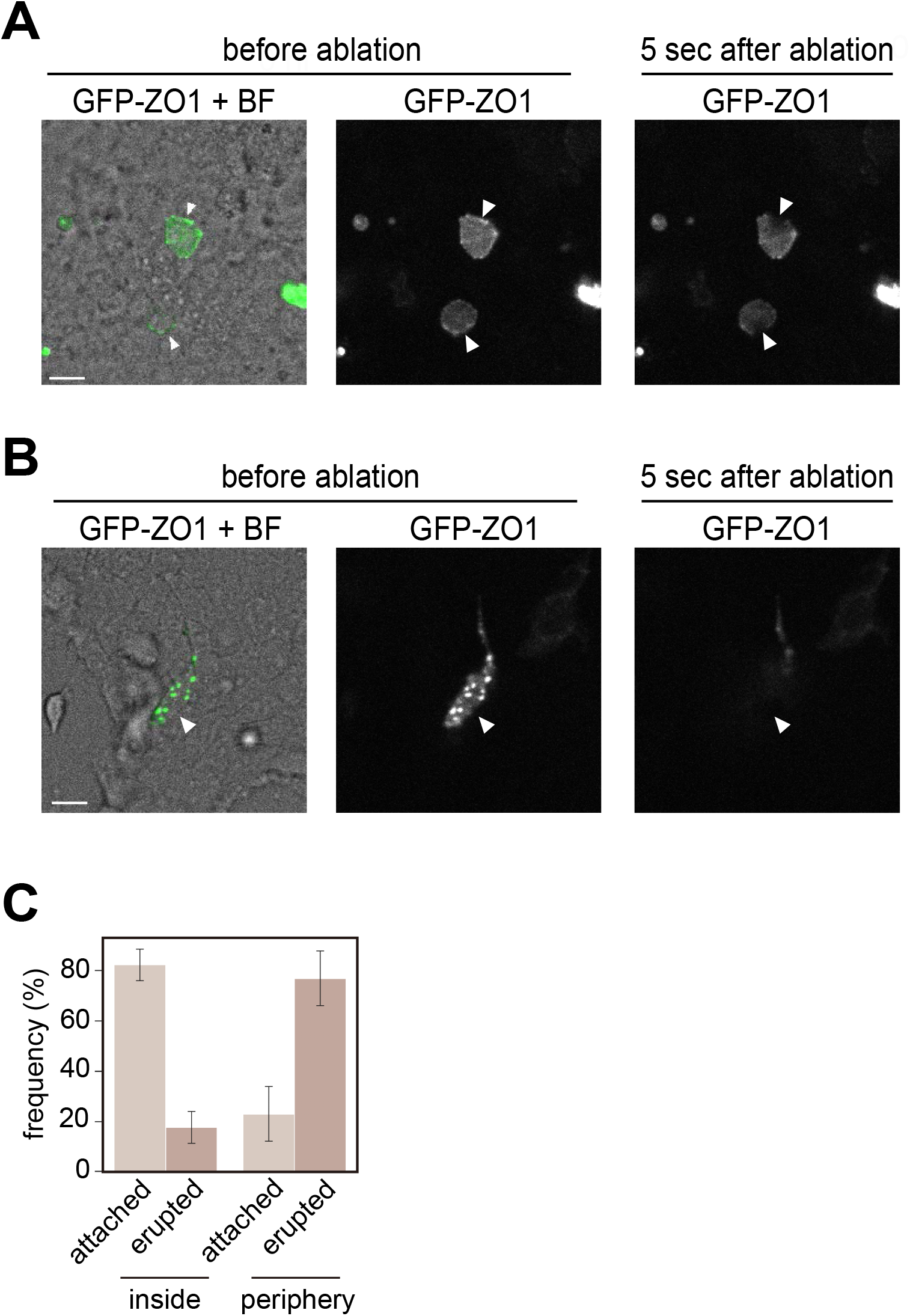
Laser ablation of MDCK cells expressing GFP-ZO-1. **A**. Cells inside an island with GFP-ZO-1 around the cell membrane. About 80% of these cells remained stable after ablation (Video S7). **B**. A cell located close to the edge of an island had GFP-ZO-1 puncta. About 80% of these cells rapidly erupted and detached from the bottom of the dish (Video S8). Arrowheads indicate laser ablation sites. **C**. A statistical summary of the laser ablation experiment. “attached” indicates cells that stayed attached to the dish after ablation, as shown in **A**. “erupted” indicates cells that erupted after ablation and detached from the dish, as shown in **B**. Data were obtained from two experiments. The total number of cells for the laser ablation were 26 inside and 24 outside cells. Scale bars, 20 μm.

**Figure S3.**
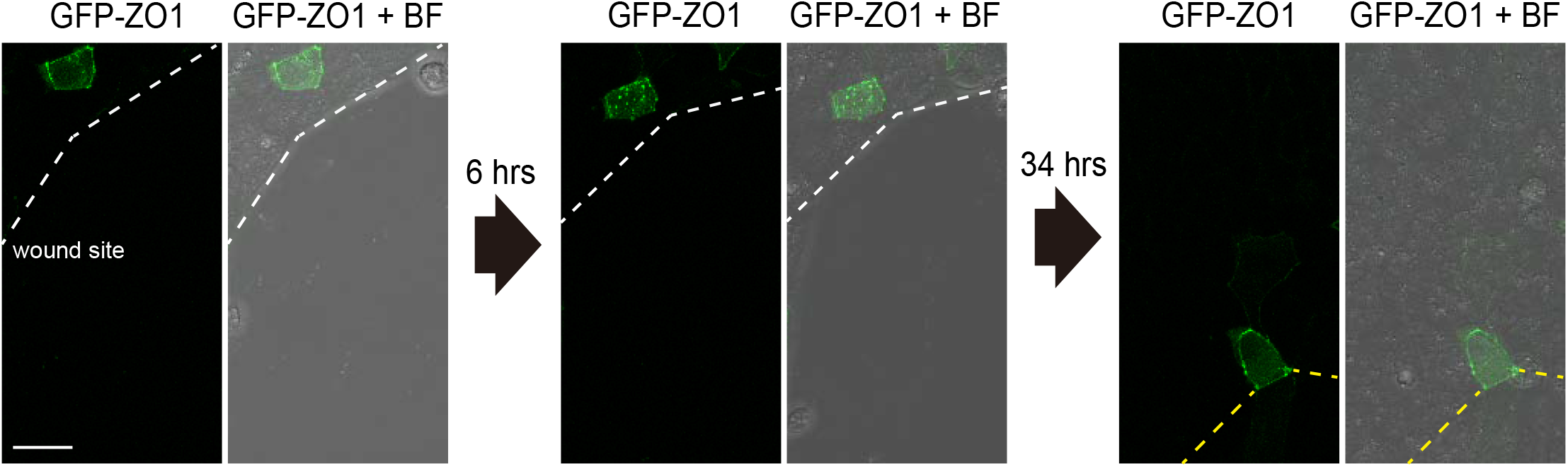
Shuttling GFP-ZO-1 between droplets and cell junction. In the wound healing assay, GFP-ZO-1 was initially localized at the cell junction (left). When cells started the migration, GFP-ZO1 formed droplets (middle). When the wound gap was filled, and a new cell-cell interaction was established, GFP-ZO-1 was re-localized to the cell junction (right). White lines indicate the wound site and yellow lines indicate the new cell-cell contact site. BF: bright field. Scale bar, 20 μm.

**Figure S4.**
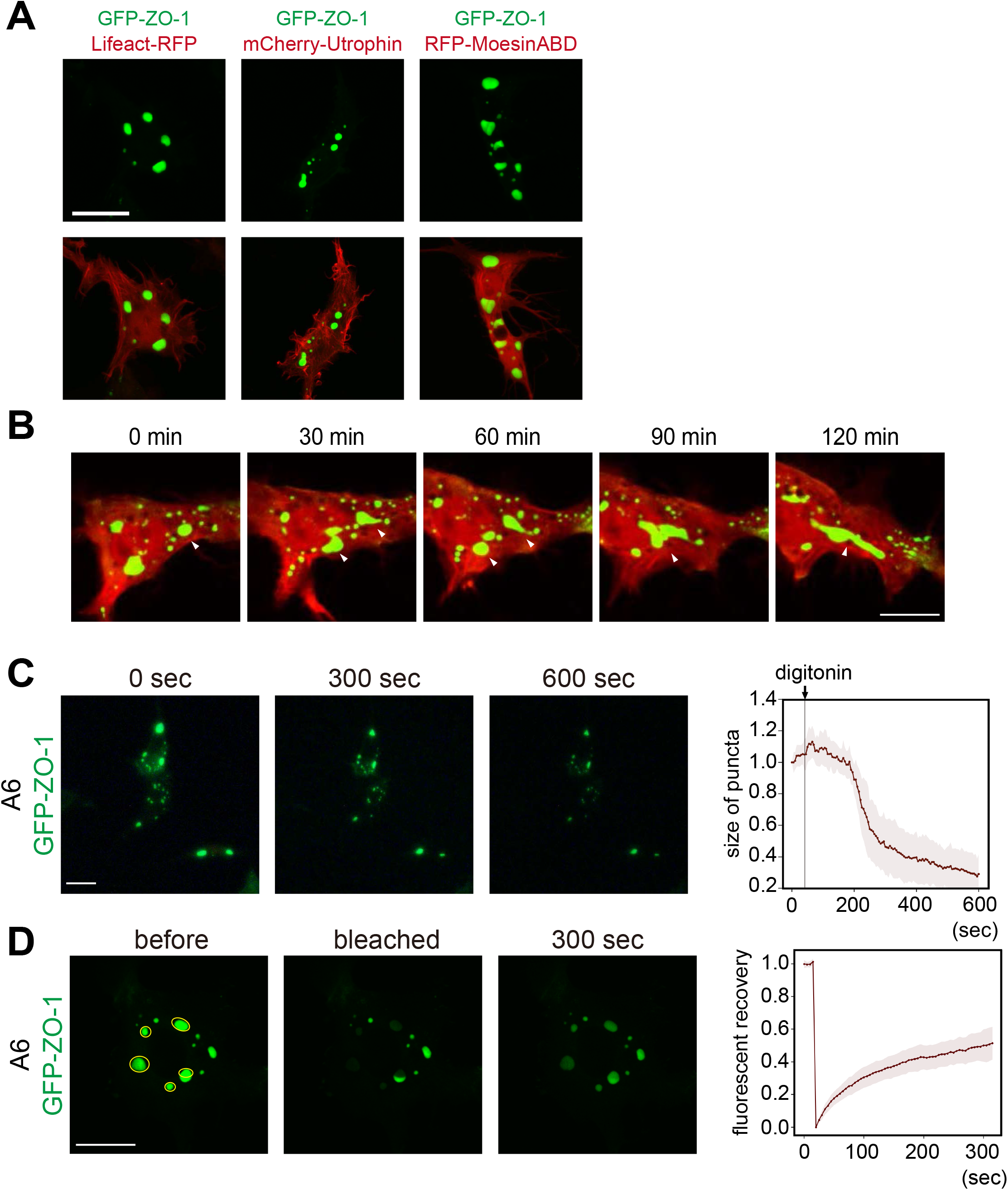
Dynamics of GFP-ZO-1 in A6 cells. **A**. GFP-ZO-1 was co-expressed with Lifeact-RFP, mCherry-Utrophin, and RFP-MoesinABD (actin-binding domain). **B**. Dynamics of GFP-ZO-1 droplets induced by co-expression with MoesinABD. Droplets often underwent fission and fusion, as indicated by arrowheads. These images are from Video S3. **C**. A6 cells expressing GFP-ZO-1 were semi-permeabilized with 0.012% digitonin. The size of puncta was quantified with Image J. **D**. Fluorescent recovery after a photobleaching (FRAP) assay of GFP-ZO-1 droplets. n = 20 puncta. Scale bars, 20 μm.

**Figure S5.**
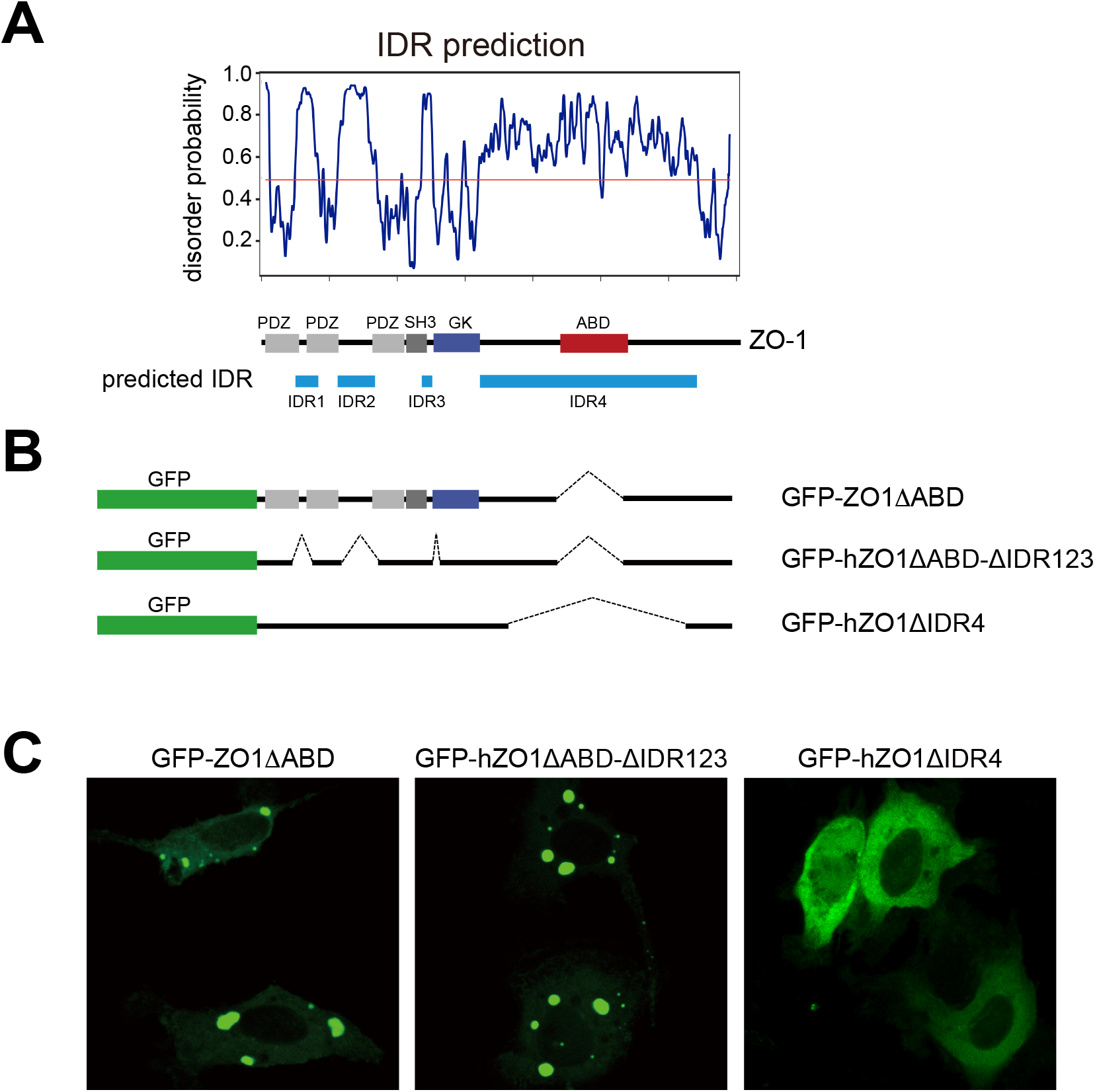
Dynamics of GFP-ZO-1 in A6 cells. **A**. IDRs (intrinsically disordered regions) of ZO-1 were predicted by the Protein Disorder Prediction System (http://prodos.hgc.jp/cgi-bin/top.cgi). Four IDRs were designated as IDR1-4. **B**. IDR deletion mutants were constructed. **C**. These mutants were expressed in A6 cells. Scale bar, 20 μm.

**Video S1.** Mouse embryos expressing ZO-1-EGFP were flushed from uteri at one-day post-coitum and observed by time-lapse microscopy for 92 h (related to Figure S1).

**Video S2**. Mouse E4.5 embryos expressing ZO-1-EGFP were pierced by a glass needle and observed by time-lapse microscopy (related to Figure 2F).

**Video S3.** An MDCK cell expressing GFP-ZO-1 was treated with digitonin (“+digitonin” at 50 sec in the video) and imaged by time-lapse microscopy (related to Figure 4D).

**Video S4.** MDCK cells expressing GFP-ZO-1 were treated with a trypsin-EDTA solution. The video starts immediately after treatment (related to Figure S4).

**Video S5.** Laser ablation of MDCK cells expressing GFP-ZO-1 inside the island. The laser was irradiated at 4 sec at the membrane indicated by arrowheads (related to Figure 5E).

**Video S6.** Laser ablation of MDCK cells expressing GFP-ZO-1 at the periphery of the island. The laser was irradiated at 4 sec at the membrane indicated by the arrowhead. The cell erupted and disappeared immediately after irradiation (related to Figure 5F).

**Video S7.** A wound-healing assay of MDCK cells. A sheet of MDCK cells transfected with the GFP-ZO-1 construct was scratched and observed by time-lapse microscopy (related to Figure 6).

**Video S8.** Shuttling GFP-ZO-1 between droplets and cell junction. In the wound healing assay, GFP-ZO-1 was initially localized at the cell junction. When cells started the migration, GFP-ZO1 formed droplets. When the wound gap was filled, and new cell-cell interaction was established, GFP-ZO-1 was re-localized to the cell junction.

**Video S9.** A wound-healing assay of MDCK cells, followed by a FRAP assay (1). A sheet of MDCK cells transfected with the GFP-ZO-1 construct was scratched and observed by time-lapse microscopy (related to Figure S5A). The cell for the following FRAP assay was indicated by an arrowhead at the first and last frames.

**Video S10.** A wound-healing assay of MDCK cells, followed by a FRAP assay (2). The FRAP assay was conducted using a cell in Video S11, which formed ZO-1 puncta (related to Figure S5B). Arrowhead indicates the granule bleached in the FRAP assay.

**Video S11.** Latrunculin B treatment of an A6 cell co-expressing GFP-ZO-1 and RFP-MoesinABD (related to Figure 3F).

**Video S12.** An A6 cell co-expressing GFP-ZO-1 and RFP-MoesinABD was imaged by time-lapse microscopy for 885 min (related to Figure S3).

**Video S13.** A6 cells co-expressing GFP-ZO-1 and RFP-MoesinABD were treated with digitonin (“+digitonin” at 50 sec in the video) and imaged by time-lapse microscopy (related to Figure 4C).

## STAR★Methods

### KEY RESOURCES TABLE

| REAGENT or RESOURCE | SOURCE | IDENTIFIER |
| --- | --- | --- |
| Antibodies |  |  |
| anti-ZO-1 antibody | Thermo Fisher Scientific | cat. # 33-9100;<br>RRID:AB_2533147 |
| goat anti-mouse IgG antibody, Alexa Fluor 488 | Thermo Fisher Scientific | cat. # A28175;<br>RRID:AB_2536161 |
| Chemicals, Peptides, and Recombinant Proteins |  |  |
| Blocking One | Nacalai Tesque | cat.# 03953 |
| KSOM | Millipore | cat. # MR-106D |
| ouabain | Sigma | cat.# O3125 |
| Alexa Fluor 546 Phalloidin | Thermo Fisher Scientific | cat.# A22283;<br>RRID:AB_2632953 |
| Hoechst 33342 | Thermo Fisher Scientific | cat.# H3570 |
| Lipofectamine 2000 | Invitrogen | cat. # 11668-027 |
| Effectene | Qiagen | cat. # 301425 |
| Latrunculin B | Enzo Life Science | cat. # BML-T110-0001 |
| digitonin | WAKO | cat. # 048-02124 |
| trypsin | Nacalai Tesque | cat # 35555-54 |
| 3-aminopropyl) triethoxysilane | Sigma-Aldrich | cat. # 440140 |
| Sulfo-SANPAH | Pierce | cat. # 22589 |
| 1,6-hexanediol | WAKO | cat. # 087-00432 |
| Experimental Models: Organisms/Strains |  |  |
| *Xenopus laevis* (J-strain) | N/A | N/A |
| R26-ZO-1EGFP | RIKEN | accession # CDB026K |
| wild type ICR mice | Japan CREA | N/A |
| Experimental Models: Cell Lines |  |  |
| MDCK I cells | Fujiwara et al. (2016) | N/A |
| A6 cells | Mimori-Kiyosue et al. (2007) | N/A |

| Recombinant DNA |  |  |
| --- | --- | --- |
| human ZO-1 | Konno et al. (2008) | N/A |
| GFP-ZO-1/pCS2 | This paper | N/A |
| GFP-ZO-1ΔABD/pCS2 (Δ1151-1371) | This paper | N/A |
| GFP-ZO-1ΔABD-ΔIDR123/pCS2<br>(Δ114-118, Δ275-415, Δ588-627,<br>Δ1151-1371) | This paper | N/A |
| GFP-ZO-1ΔIDR4/pCS2 (Δ858-1580) | This paper | N/A |
| GFP-ZO-1-PSG/pCS2 (424-794) | This paper | N/A |
| GFP-ZO-1ΔABDΔPSG/pCS2 (Δ424--794,<br>Δ1151-1371) | This paper | N/A |
| Lifeact-RFP | Iioka et al. (2007) | N/A |
| RFP-MoesinABD | Iioka et al. (2004) | N/A |
| mCherry-Utrophin | Kim and Davidson (2013) | N/A |
| Software and Algorithms |  |  |
| Fiji | NIH | https://fiji.sc |
| Python 3.6 | Python Core Team | https://www.python.org |
| Pandas | v0.22 | http://pandas.pydata.org |
| seaborn | v0.8 | https://seaborn.pydata.org |
| Matplotlib | v2.2 | https://matplotlib.org |
| Andor IQ2 | Oxford Instruments | https://andor.oxinst.com/products/iq-live-cell-imaging-software/ |
| PrDOS (protein disorder prediction system) | (Ishida and Kinoshita, 2007) | http://prdos.hgc.jp/cgi-bin/top.cgi |

#### Lead Contact and Materials Availability

Further information and requests for resources and reagents should be directed to and will be fulfilled by the Lead Contact, Naoto Ueno. All unique/stable reagents generated in this study are available from the Lead Contact with a completed Material Transfer Agreement.

#### Method details

##### Mouse embryo collection

Animal care and experiments were conducted in accordance with the Guidelines of Animal Experimentation of the National Institutes for Natural Sciences. All animal experiments were approved by the Animal Research Committee of the National Institutes for Natural Sciences. Mice were maintained in a light- and temperature-controlled room using a 12 h light:12 h dark cycle at 23+/-2 °C.

Males of R26-ZO-1EGFP (Katsunuma et al., 2016) or wild-type ICR (Japan CREA) were mated with wild-type ICR females to obtain blastocysts. Preimplantation E3.5 embryos were flushed from uteri with KSOM (MR-106D, Millipore). Embryos were cultured in KSOM covered with mineral oil at 37 °C and 5% CO_2_. Fluorescent signals were monitored using a confocal microscope (A1, Nikon) or a spinning disc confocal microscope (CV1000, Yokogawa, Japan). For immunofluorescence, embryos were fixed at 4 °C in 4% paraformaldehyde (PFA) in PBS overnight. For ouabain treatment, 10 mM ouabain (cat. # O3125, Sigma) in DMSO was diluted to the indicated concentrations.

##### Manipulation of mouse embryos

To remove the zona pellucida, embryos were treated with Tyrode’s solution (Bradley, 1987). Embryos were transferred to a drop of Tyrode’s solution. The zona was removed typically within 1 min. Then, embryos were washed three times in KSOM. To pierce the mural trophectoderm, microinjection needles were made from a 1 mm diameter glass capillary (GD1, Narishige, Japan), the needle was set to a micromanipulator (Leica), and embryos were pierced with the needle by manual manipulation.

For hexanediol treatment, 1.6-hexanediol (cat. # 087-00432, WAKO, Japan) was dissolved in water at 50% w/v, and a 1/10 volume was added to the culture medium. Fluorescent images were captured using CV1000 (Yokogawa, Japan).

##### Immunofluorescence with mouse embryos

Fixed embryos were washed three times with 1% bovine serum albumin (BSA) in PBS and then soaked in a blocking solution, 0.1% Triton X-100 in Blocking One (cat. # 03953, Nacalai, Japan), for 1 h at room temperature. Embryos were then incubated in primary antibody solution containing an anti-ZO-1 antibody (cat. # 33-9100, Thermo) diluted 100-times in Blocking One at 4 °C overnight. After washing three times with the blocking solution, embryos were incubated in a secondary antibody solution containing 100-times diluted goat anti-mouse IgG antibody Alexa Flour 488 (cat. # A27185, Thermo), 100-times diluted Alexa Fluor 546 Phalloidin (cat. # A22283, Thermo Fisher), and 10 μg/ml Hoechst 33342 (cat. # H3570, Thermo Fisher) in Blocking One. Fluorescent signals were monitored using Nikon A1 or Leica SP8 confocal microscopes.

##### Measurement of fluorescent intensities and particle analyses

Fiji/Image J was used to measure fluorescent intensities. Particle analyses in Fiji/Image J were used to count particles and measure particle areas. A large mass signal in the cavity was removed for quantitative analyses. Images were inverted and converted to an 8-bit type. Cytoplasmic regions were manually determined. The “Analyze Particles” program was run with the threshold signal value: 230, circularity: 0.3-1.0, and size: 3 - infinity.

##### GFP-ZO-1 expression constructs

Human ZO-1 cDNA was a gift from Dr. Fumio Matsuzaki (Konno et al., 2008). EGFP-human ZO-1/pCS2 was constructed by Dr. Makoto Suzuki. ZO-1ΔABD lacks amino acid (aa) # 1151-1371. Lifeact-RFP is described in Iioka et al. (2007). MoesinABD is described in Iioka et al. (2004). The mCherry-Utrophin construct was a gift from Lance A. Davidson (Kim and Davidson, 2013).

##### Cell culture and transfection of plasmids

A6 cells, established from a normal *X. laevis* kidney, were a gift from Dr. Yuko Mimori-Kiyosue (Mimori-Kiyosue et al., 2007). A6 cells were grown at 24 °C without CO2 in Leibovitz’s L-15 medium (50% L-15 medium, 10% fetal bovine serum (FBS), 200 mg/l kanamycin). MDCK cells were a gift from Dr. Mitsuru Nishita and Dr. Kensaku Mizuno. MDCK cells were cultured in Dulbecco’s modified Eagle medium (DMEM) containing 10% FBS. Transfection was conducted using Lipofectamine 2000 (Invitrogen # 11668-027) for MDCK cells and Effectene (Qiagen # 301425) for A6 cells following the manufacturers’ instructions.

##### Immunofluorescence of the tissue culture cells

Cells were fixed in 4% PFA in PBS at 4 °C overnight, washed with 0.1% Triton X-100 in PBS. Blocking One (cat. # 03953, Nacalai Tesque, Japan) was used to block for 1 h at room temperature, followed by incubation in the primary antibody solution, a 100-times diluted anti-ZO-1 antibody (cat. # 33-9100, Thermo) in Blocking One at 4 °C overnight. After washing three times in Blocking One, cells were incubated in the secondary antibody solution containing 200-times diluted goat anti-mouse IgG antibody Alexa Fluor 488 (cat. # A27185, Thermo Fisher) and 200-times diluted Alexa Fluor 546 Phalloidin (cat. # A22283, Thermo Fisher) in Blocking One at 4 °C overnight. After washing three times in PBS, cells were observed using a Leica SP8 confocal microscope.

##### Laser ablation

Laser ablation was conducted using an Olympus IX 81 inverted microscope (20 x /0.70 NA dry objective lens), equipped with a spinning-disc confocal unit Yokogawa CSUX-1 and iXon3 897 EM-CCD camera (Andor), controlled with Andor IQ2 software. An N2 Micropoint laser (16 Hz, 365 nm, 2.0 μW, Photonic Instruments) was focused on the membrane at the cell membrane. Time-lapse images were acquired every 200 msec before and during the laser ablation process and analyzed with Fiji/ImageJ software.

##### Fluorescence recovery after photobleaching (FRAP)

FRAP assays in the tissue culture cells were conducted with a Leica SP8 confocal microscope and software equipped in the microscope operating system. Regions of interest (ROI) were bleached using a 488 nm laser. Pre-bleach and post-bleach images were acquired with a 488 nm laser. Fluorescence recovery of GFP-ZO-1 was monitored. Recovery data were background corrected and normalized to the ROI intensity before bleaching. A reference ROI outside the bleached area was processed in the same way.

##### Observation of GFP-ZO-1 dynamics in the tissue culture cells

For the wound-healing assay, MDCK cells almost confluently grown on a glass-bottom dish were transfected with GFP-ZO-1/pCS2. After one day, the cell layer was scratched with a Pipetman tip, incubated at 37 °C, 5% CO_2_ in a stage-top incubator (cat. # STXG, Tokai HIT, Japan), and observed by a Leica SP8 confocal microscope.

For the latrunculin B treatment, 2.5 μM latrunculin B in the culture medium was added to the culture dish of A6 cells transiently expressing GFP-ZO-1 so that the final concentration of latrunculin B was 0.5 μM. For the digitonin treatment, 0.06% digitonin in the culture medium was added to the culture dish so that the final concentration of digitonin was 0.012%. For the hyperosmolarity treatment of A6 cells, 10 × PBS was added so that the concentration of PBS changed from 0.5 × to 1.5 ×. Cells were observed at room temperature. For the trypsin-EDTA treatment, the culture medium of MDCK cells was replaced with PBS containing 0.05% trypsin and 1 mM EDTA.

##### Preparation of polyacrylamide-based plates

Glass bottom dishes (27-mm diameter, cat.# 3910-035, IWAKI, Japan) were treated with 2% (v/v) (3-aminopropyl) triethoxysilane (cat.# 440140 Sigma-Aldrich)/acetone. Air-dried plates were washed with sterilized water three times and dried at 50 °C for 2 h. 45 μl acrylamide solution was loaded, and then a 24-mm diameter coverslip was loaded onto the solution. After gelation, the coverslip was removed, and the surface of gelled polyacrylamide was washed with 50 mM HEPES (pH 8.5). 200 μl of 0.5 mg/ml Sulfo-SANPAH (cat.# 22589, Pierce) was loaded and irradiated with UV for 30 min using Stratalinker 2400 (Stratagene). Then, it was washed twice with 50 mM HEPES (pH 8.5), and 200 μl of 100 μg/ml fibronectin was loaded and incubated overnight at 4 °C. It was washed with the MDCK culture medium to block residual reactive groups, and cells were plated.

##### Atomic force microscopy (AFM) measurements

AFM measurements were conducted with a JPK Nanowizard Cellhesion 200 (JPK Instruments AG, Germany) fitted to an Axio Zoom V. 16 system (Zeiss). For a cantilever, CP-CONT-BSG-A (sQUBE, Germany) was used. Young’s modulus was calculated using JPK data processing software (Bruker). Measurement conditions were as follows: setpoint 50-200 nN, approach speed 2 μm/s, data rate 1000 Hz, measured spring constant 0.342 N/m.

